# Structural basis of the allosteric regulation of cyanobacterial glucose-6-phosphate dehydrogenase by the redox sensor OpcA

**DOI:** 10.1101/2024.02.29.582749

**Authors:** Sofía Doello, Dmitry Shvarev, Marius Theune, Jakob Sauerwein, Alexander Klon, Erva Keskin, Marko Boehm, Kirstin Gutekunst, Karl Forchhammer

**Author notes:** SD and KF conceptualized the study. SD, MT, JS, AK, EK, and MB carried out the experiments. DS acquired and analyzed cryo-EM data. SD and DS wrote the manuscript. All authors participated in data interpretation and manuscript edition. The authors declare no competing interests. S.D. contributed equally to this work with D.S.

## Abstract

The oxidative pentose phosphate (OPP) pathway is a fundamental carbon catabolic route for generating reducing power and metabolic intermediates for biosynthetic processes. In addition, its first two reactions form the OPP shunt, which replenishes the Calvin-Bassham cycle under certain conditions. Glucose-6-phosphate dehydrogenase (G6PDH) catalyzes the first and rate-limiting reaction of this metabolic route. In photosynthetic organisms, G6PDH is redox-regulated to allow fine-tuning and to prevent futile cycles while carbon is being fixed. In cyanobacteria, regulation of G6PDH requires the redox protein OpcA, but the underlying molecular mechanisms behind this allosteric activation remain elusive. Here, we used enzymatic assays and *in vivo* interaction analyses to show that OpcA binds G6PDH under different environmental conditions. However, complex formation enhances G6PDH activity when OpcA is oxidized and inhibits it when OpcA is reduced. To understand the molecular basis of this regulation, we used cryogenic electron microscopy to determine the structure of cyanobacterial G6PDH and the G6PDH-OpcA complex. OpcA binds the G6PDH tetramer and induces conformational changes in the active site of G6PDH. The redox sensitivity of OpcA is achieved by intramolecular disulfide bridge formation, which influences the allosteric regulation of G6PDH. *In vitro* assays reveal that the level of G6PDH activation depends on the number of bound OpcA molecules, which implies that this mechanism allows delicate fine-tuning. Our findings unveil a novel and unique molecular mechanism governing the regulation of the OPP pathway in cyanobacteria.

---

Glucose-6-phosphate dehydrogenase (G6PDH) catalyzes the first and rate-limiting step of the oxidative pentose phosphate pathway (OPP), converting glucose-6-phosphate to 6-phosphogluconolactone (Figure 1A). This pathway is essential for nucleotide synthesis and plays an important role in balancing the cellular redox state by controlling the NADPH/NADP^+^ ratios. Therefore, regulation of G6PDH activity is of key importance in all organisms (1).

**Fig. 1.**
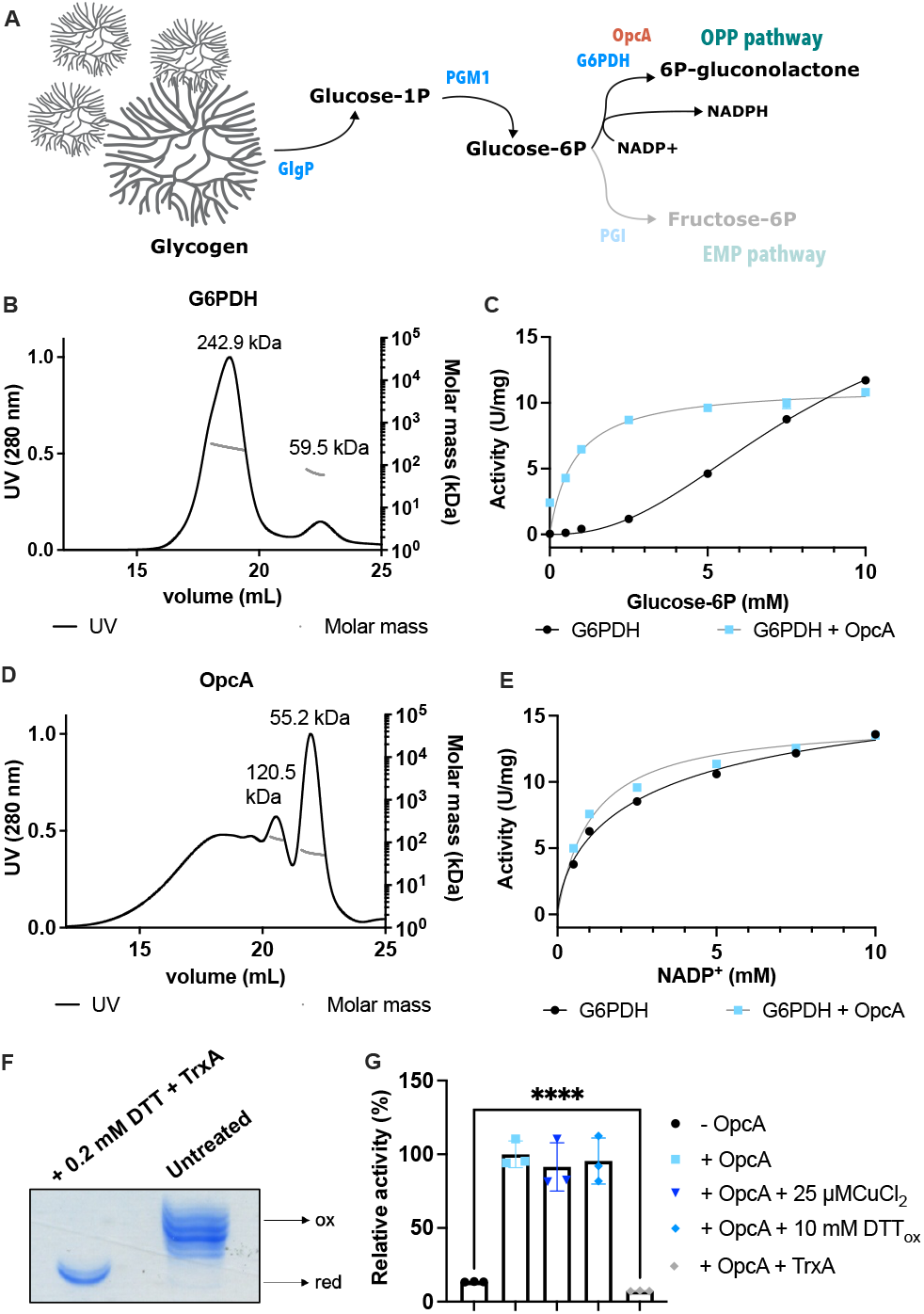
Effect of OpcA on G6PDH activity. (A) Scheme of the role of G6PDH in carbon catabolism. (B) SEC-MALS analysis of recombinant G6PDH. (C) Activity of recombinant G6PDH in the absence (black) and presence (blue) of OpcA. (D) SEC-MALS analysis of recombinant OpcA. (E) Activity of recombinant G6PDH with a fixed glucose-6-P concentration of 10 mM and increasing concentrations of NADP^+^ in the absence (black) and presence (blue) of OpcA. (F) PEGylation assay of recombinant OpcA before and after incubation with 0.2 mM DTT and TrxA. (G) Activity of recombinant G6PDH in the absence (black) and presence of untreated OpcA (blue) and OpcA pre-treated with different redox agents (shades of blue and gray). Every condition was measured at least three times. Error bars represent the standard deviation (SD). Error bars are not visible only when they are smaller than the icon size. Four asterisks represent a p-value ≤ 0.0001.

In mammals, G6PDH is of special relevance in erythrocytes, which lack mitochondria and rely on the pentose phosphate pathway to produce NADPH (2). In photosynthetic organisms, G6PDH is especially needed during the night and certain nutritional conditions (e.g. resuscitation from nitrogen starvation), when glucose catabolism is the sole source of energy and metabolites (3). Thus, G6PDH is mostly active under such environmental contexts and less active during photoautotrophic growth. In chloroplasts, this regulation is achieved via redox control. When photosynthesis is operating, G6PDH is kept in a reduced state by the thioredoxin system. In the dark, G6PDH is oxidized and disulfide bond formation leads to activation of the enzyme (4). In addition to its role as a key enzyme of a carbon catabolic pathway in the dark, G6PDH is part of the so-called glucose-6-phosphate shunt in plant chloroplasts and the OPP shunt in cyanobacteria, both of which designate a bypass starting with glucose-6-phosphate that feeds carbon into the Calvin-Benson cycle in the form of ribulose-5-phosphate (5, 6). These shunts enhance the metabolic flexibility and stabilize photosynthesis, especially during transition states and under changing redox conditions. Therefore, the activity of plastidic and cyanobacterial G6PDH requires a delicate mode of regulation. For instance, *Arabidopsis thaliana* possesses three functional plastidic G6PDH isoforms which are differently regulated and partly active in the light, which was suggested to allow flux via the glucose-6-phosphate shunt into the Calvin-Benson cycle while simultaneously minimizing futile cycling (4). In cyanobacteria, the only prokaryotes capable of carrying out oxygenic photosynthesis, the redox-sensitive cysteine residues present in plastidic plant G6PDH are not conserved (Figure S1A). Redox regulation of cyanobacterial G6PDH is achieved through an auxiliary protein, termed oxidative pentose phosphate cycle protein (OpcA) (7).

Although OpcA has previously been reported as a unique cyanobacterial protein, a homologue with significant similarity is present in actinobacteria (Figure S1B). Among cyanobacteria, OpcA is highly conserved. This protein was initially suggested to act as a chaperone for the assembly of G6PDH oligomers (8), but it was later shown to rather act as an allosteric activator of G6PDH (7). The redox state of OpcA is controlled by m-type thioredoxins, and when OpcA is oxidized, it activates G6PDH (9). However, the structural mechanism of this allosteric activation was so far unknown.

Here, we used the unicellular cyanobacterial strain *Synechocystis* sp. PCC 6803 (hereafter *Synechocystis*) to elucidate the mechanism of activation of G6PDH by OpcA and to study the structural dynamics of the complex. By enzymatic assays we demonstrate that G6PDH activity is both positively and negatively regulated by the binding of OpcA, depending on the redox state of the latter. Our high-resolution structures reveal that OpcA forms a complex with the G6PDH tetramer and induces conformational rearrangements in the active site of G6PDH. The redox sensitivity of OpcA is explained by the importance of disulfide bridge formation within OpcA for the allosteric regulation of G6PDH activity. The G6PDH-OpcA complex is, however, present under oxidative and reduced conditions *in vivo*. Altogether, our findings reveal a novel and unique molecular mechanism for OPP pathway regulation in cyanobacteria.

## Results

### OpcA acts as an activator of G6PDH depending on its redox state

To confirm an allosteric activating effect of OpcA on *Synechocystis* G6PDH, a His-tagged version of each protein was recombinantly produced in *Escherichia coli*. Size-exclusion chromatography coupled with multi-angle light scattering analysis (SEC-MALS) revealed that the most abundant oligomeric state of G6PDH has a mass of 242.9 kDa (Figure 1B), which likely represents a homotetramer. Measurement of the *in vitro* activity of G6PDH showed a sigmoidal kinetic profile with a Hill coefficient of 2.3 (Figure 1C), which indicates positive cooperative binding of glucose-6-phosphate to the G6PDH tetramer. OpcA was mostly present as a monomer, although higher oligomeric states were also detected by SEC-MALS analysis (Figure 1D). The presence of OpcA dramatically changed the kinetic profile of the G6PDH reaction: instead of the sigmoidal profile with half maximal velocity at 8.09 mM, G6PDH now displayed a Michaelis-Menten kinetic profile with highly increased substrate affinity (Figure 1C), with a a Km for glucose-6-phosphate of 0.78 mM with OpcA. The effect of OpcA was limited to glucose-6-phosphate and did not significantly affect the Km for NADP^+^ (Figure 1E). The described experiments were carried out without any redox treatment of the recombinant proteins. A PEGylation assay revealed that recombinantly purified OpcA was mostly in an oxidized state (Figure 1F). To clarify how the redox state of OpcA affects G6PDH activity, OpcA was pre-incubated with recombinant *Synechocystis* TrxA and 0.2 mM DTT for 30 min to achieve full reduction (Figure 1F). OpcA only showed activation of G6PDH when it was in an oxidized state, whereas the activity of G6PDH was significantly lower in the presence of reduced OpcA (Figure 1G), suggesting that OpcA can act as an activator of G6PDH when it is oxidized, and as an inhibitor when it is reduced. As expected, treatment of OpcA with oxidizing agents (DTTox or CuCl_2_) did not significantly affect G6PDH activity since untreated OpcA was already in an oxidized state.

### OpcA interacts with the allosteric site of G6PDH and stabilizes its N-terminus

To uncover how OpcA regulates G6PDH, we used cryogenic electron microscopy (cryo-EM) to determine the structure of both G6PDH and the G6PDH-OpcA complex. Cyanobacterial proteins were recombinantly produced and purified using affinity chromatography. We then performed sample quality control using negative-stain electron microscopy (EM) (Figure S2A). Our further single-particle cryo-EM analysis of the G6PDH-OpcA sample revealed a homogeneous distribution of particles. This allowed us to generate 2D class averages that display high-resolution structural features (Figure S2B). Using a total of 6351 micrographs, we performed multiple rounds of 2D and 3D classifications in cryoSPARC (10) to refine the structures of the G6PDH tetramer and tetrameric G6PDH bound to OpcA (Figure 2, Movies S1,2). The resolutions achieved were 3.3 Å (269214 particles) and 3.7 Å (128652 particles) respectively (Figures S2C,D, S3). Additionally, we detected minor amounts of other oligomeric states of G6PDH, including trimers and larger complexes of interacting tetra- and trimers. However, we did not observe monomers or dimers, as previously reported for homologous enzymes (11–14).

**Fig. 2.**
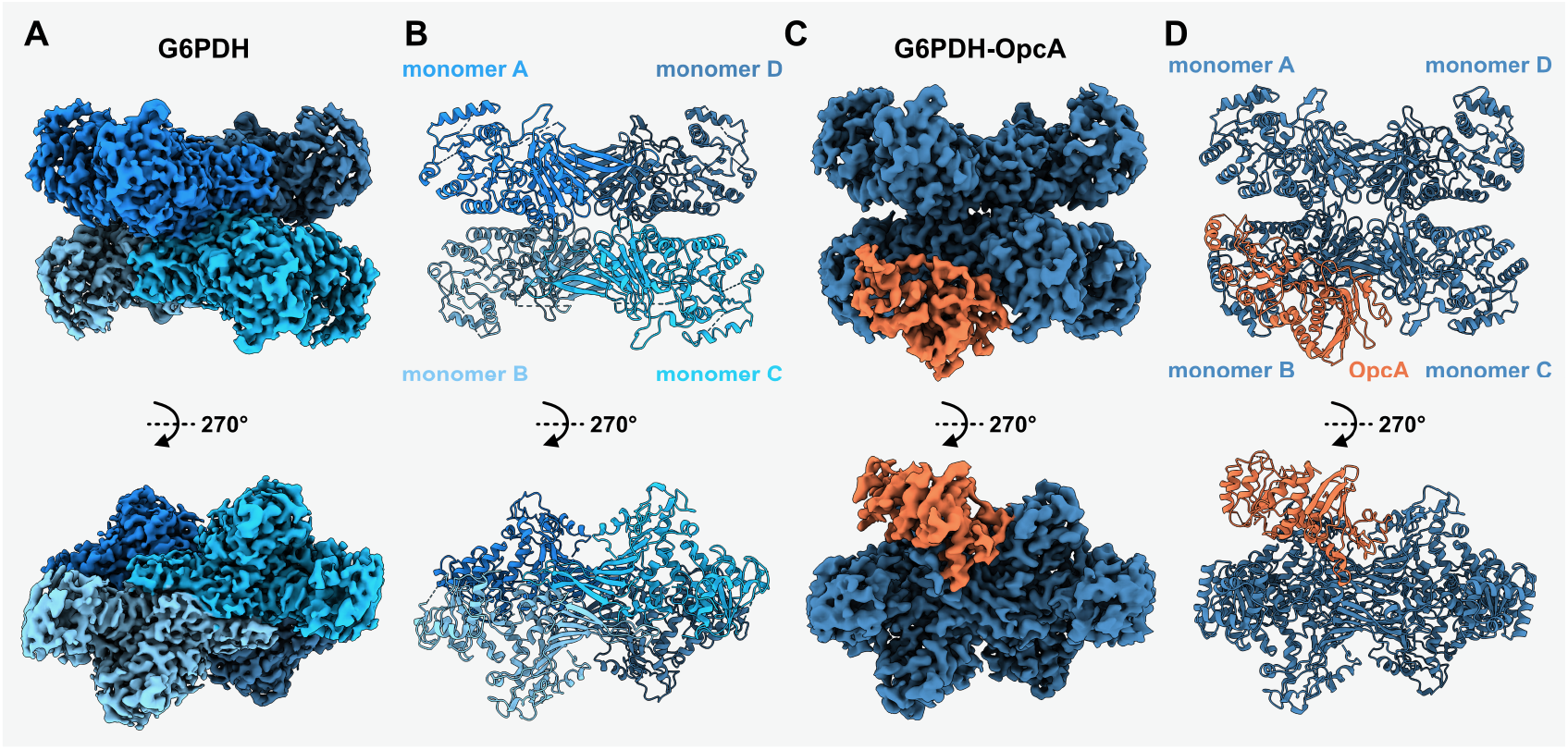
Overall architecture of cyanobacterial G6PDH and G6PDH-OpcA complex. (A) D2 symmetric cryo-EM map of the *Synechocystis* G6PDH tetramer. The map is colored by monomers, shown are the top and side views. (B) Molecular model of the *Synechocystis* G6PDH tetramer, colored and viewed as in (A). (C) Cryo-EM map of the *Synechocystis* G6PDH-OpcA complex. OpcA is colored orange, G6PDH is colored dark blue. The map is viewed as in (A). (D) Molecular model of the *Synechocystis* G6PDH-OpcA complex, colored and viewed as in (C).

The tetrameric structure of cyanobacterial G6PDH displays a D2 symmetry and consists of two G6PDH dimers (Figures 2A,B,S4). This structure is similar to the G6PDH structures found in other organisms (Figure S5). In contrast, the G6PDH-OpcA complex is not symmetric and has only one OpcA molecule bound to one of the dimers (monomers B and C) within the G6PDH tetramer (Figure 2C,D). OpcA is a relatively small protein of 52 kDa (Figure 1D) consisting of the N-terminal-, C-terminal-, and a small alpha-helical putative peptidoglycan binding domain (Figure 3A). Binding with G6PDH occurs between the OpcA C-terminal domain and the N-terminus of G6PDH monomer B and the C-terminal domain of G6PDH monomer C (Figure 3B,C). Interestingly, the OpcA putative peptidoglycan binding domain, whose position in the complex would be far from the binding interface with G6PDH, is not resolved in our structure (Figure 3A-C). The interaction of G6PDH with OpcA occurs primarily in two areas. The first interface is formed by the contacts between the structured C-terminal loops, helix *α*o and the C-terminus of G6PDH monomer C with the C-terminal region of OpcA (Figure 3D). Remarkably, the region of OpcA (T407-T434) that interacts with G6PDH monomer C contains two cysteine residues: C410 and C416 (Figure 3D). These cysteine residues form a disulfide bond in the oxidized state of the protein, which is the natural state of purified OpcA (Figure 1F). This disulfide bridging contributes to the special conformational arrangement of OpcA, which promotes the allosteric regulation of G6PDH. The second interface is created by the structured N-terminus of G6PDH monomer B (amino acid residues 5-11) binding to the *α*-helix Q298-S312 of OpcA (Figure 3E). Notably, monomer B is the only monomer in the G6PDH tetramer with the structured N-terminus, which is supported by the interaction with OpcA.

**Fig. 3.**
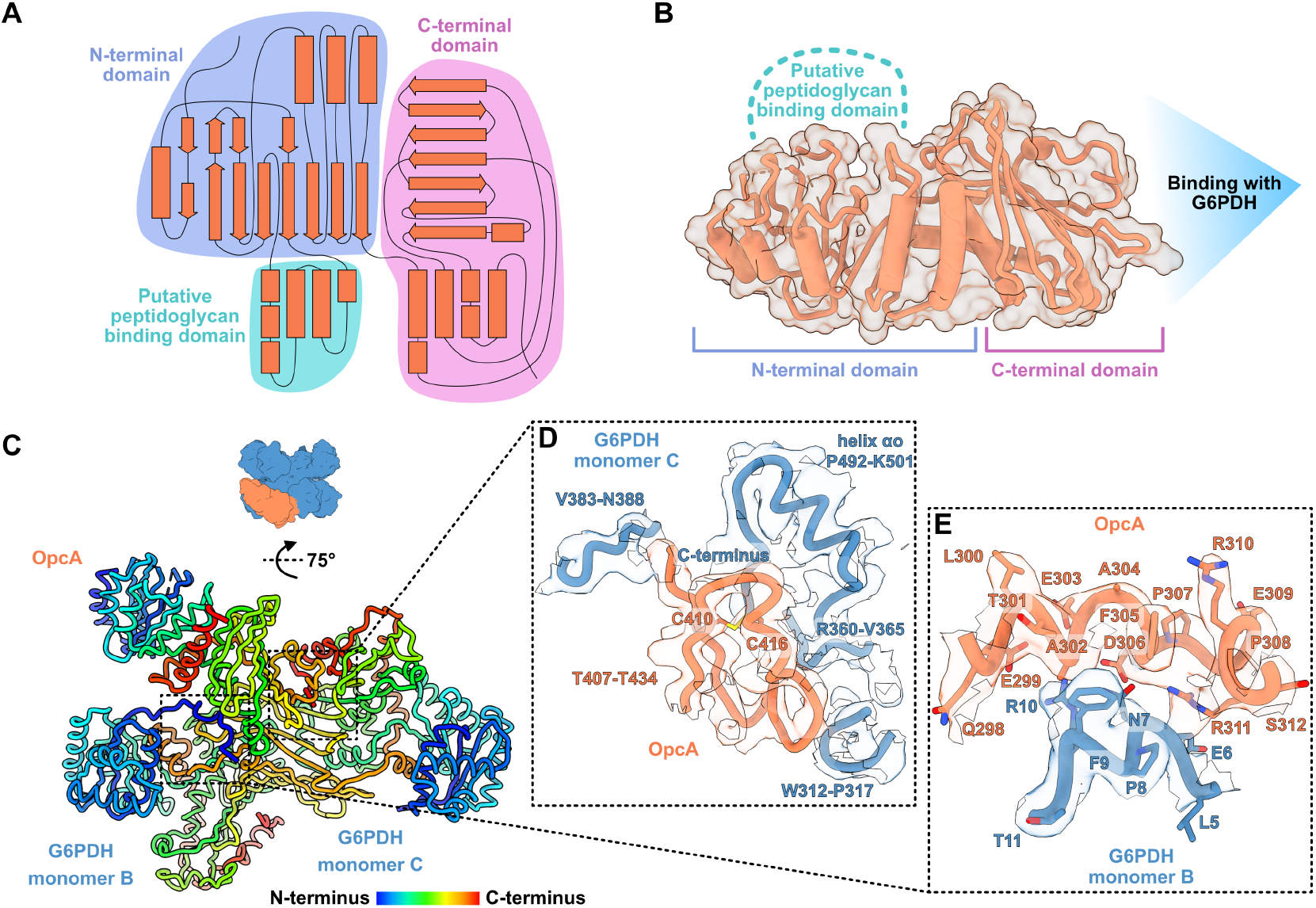
*Synechocystis* OpcA structure and interactions with G6PDH. (A) Schematic representation of the OpcA secondary structure, domains are indicated. (B) Tertiary structure of OpcA from the G6PDH-OpcA complex in cartoon (orange) and surface (semi-transparent) representation. (C) G6PDH-OpcA complex viewed from the side. Proteins are shown as ribbons colored by amino acid sequence. The model is fitted into the cryo-EM (Figure 2C) map (semi-transparent envelope). (D, E) Close-up views of the interaction interfaces between G6PDH (dark blue) and OpcA (orange).

### Structural changes in G6PDH-OpcA complex and oligomeric state analysis

At the quaternary structure level, the binding of OpcA to G6PDH does not cause global rearrangements within the G6PDH tetramer, but only slightly shifts the relative positions of the monomers (Figure S5B). However, when compared with a monomer from the G6PDH apo structure, each G6PDH monomer appeared to have undergone structural changes upon binding to OpcA (Figure 4A). Remarkably, major conformational changes were observed in the vicinity of the substrate binding sites of G6PDH monomers B and C, which directly interact with OpcA (Figure 4B-E). These included rearrangements within the conserved *α*-helix *α*f’, which is involved in the binding of the ligand glucose-6-phosphate and catalysis (14), the closely located loop and helix *α*m, as well as residues in the helix *α*a, which participates in the binding of the catalytic molecule of NADP^+^ (Figure 4B,C). In particular, the positions of residues H197 and Y198, which are essential for glucose-6-phosphate binding, are changed in both monomers B and C in the G6PDH-OpcA complex (Figure 4D-E). In contrast, the side chain orientation H259, which is essential for catalysis (15), remains unaltered upon binding of OpcA (Figure 4D-E).

**Fig. 4.**
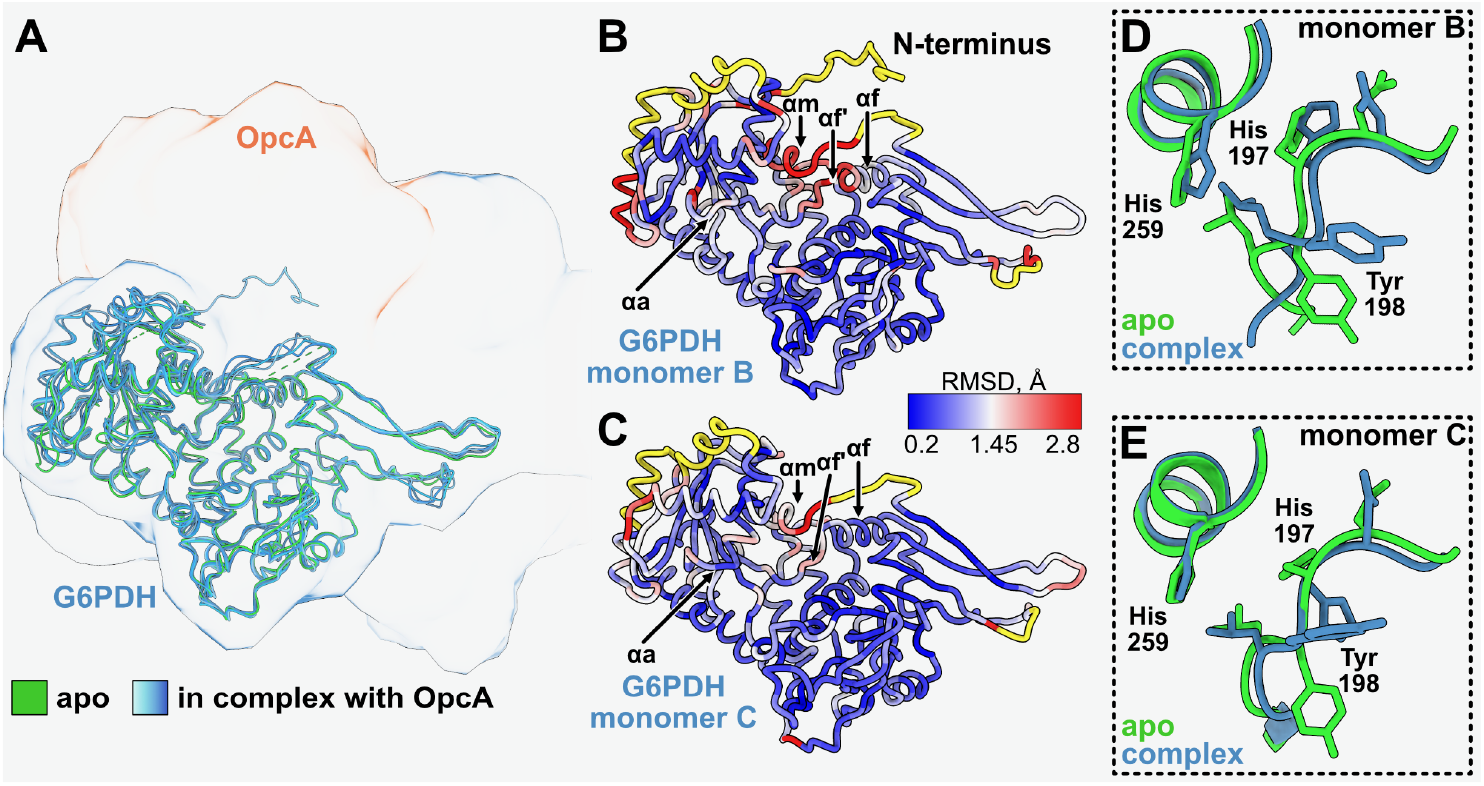
Structural changes in *Synechocystis* G6PDH induced by OpcA binding. (A) G6PDH monomers from the G6PDH-OpcA complex (different shades of blue) superimposed with a G6PDH monomer from the apo structure (green). (B, C) Structural variability of G6PDH monomer B (B) and monomer C (C) from the G6PDH-OpcA complex in comparison to the respective monomers from the G6PDH apo structure. Structures are shown in ribbon representation and are colored by the root-mean-square deviation (RMSD). Protein fragments without an RMSD value are highlighted in yellow as they are not present in the apo structure. Structured N-terminus of monomer B is indicated. (D, E) Structural changes in the active site of G6PDH monomer B (D) and monomer C (E) upon OpcA binding (blue) in comparison to the apo structures (green).

According to our cryo-EM analysis, the enzyme complex contained four subunits of G6PDH and one subunit of OpcA (Figure 2). To determine whether this is the most stable form of the complex or if other oligomeric states also exist, we first did a titration to find out at which concentration of OpcA saturation of G6PDH activation is reached. This was achieved by measuring the activity of G6PDH in the presence of different amounts of OpcA (G6PDH-OpcA molar ratios of 4:1, 4:2, 4:3, 4:4, and 4:8) (Figure 5A). The activation of G6PDH reached saturation only when OpcA was present at an equimolar concentration, suggesting more than one molecule of OpcA can bind the G6PDH tetramer. Subsequently, we analyzed the size of the complex at the described ratios using mass photometry. At a 4:1 G6PDH:OpcA ratio, a complex with a mass of 300 kDa was detected, which corresponds to a G6PDH tetramer bound to a monomeric OpcA, the same complex detected via cryo-EM (Figure 5B). However, when both proteins were present in equimolar amounts (4:4), the size of the complex increased to 450 kDa (Figure 5C), corresponding to four OpcA molecules bound to the G6PDH tetramer. When OpcA was present in excess (8:4), the size of the complex did not change (Figure 5D), suggesting that after an equimolar concentration, the interaction saturation is reached. However, analysing the relative abundance of G6PDH and OpcA during photoautotrophic growth via western blot indicated that G6PDH is more abundant than OpcA (Figure 5E), suggesting the complex is unsaturated *in vivo*.

**Fig. 5.**
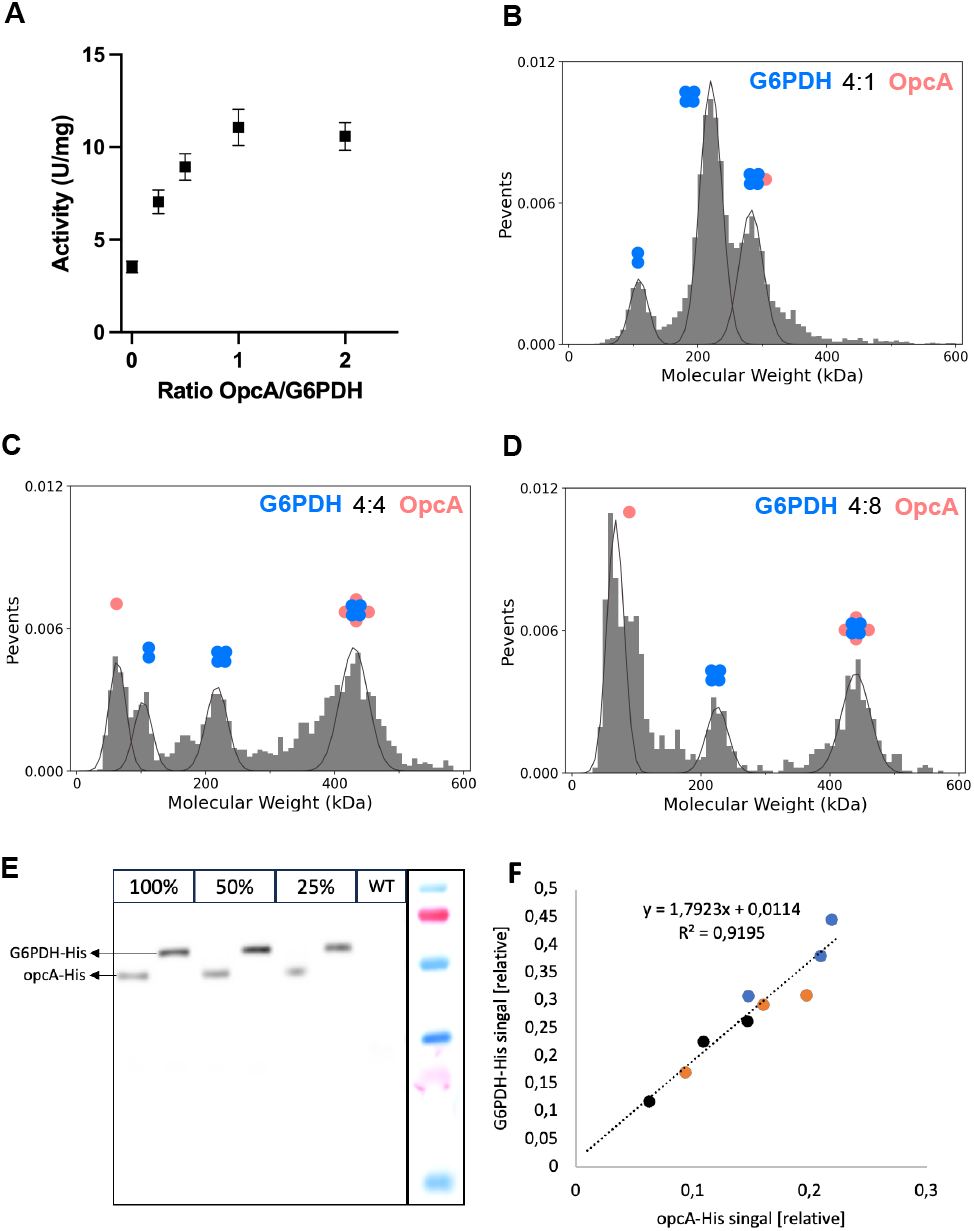
Oligomeric state analysis of the G6PDH-OpcA complex. (A) Activity of G6PDH in the presence of increasing concentrations of OpcA. Every condition was measured at least three times. Error bars represent the SD. (B-D) Mass photometry analysis of the G6PDH-OpcA complex in the presence of increasing concentrations of OpcA. (E) Western blot analysis of the relative abundance of His tagged G6PDH compared to OpcA during vegetative growth. (F) Western blot signal of G6PDH-His plotted against the signal from OpcA-His, colors indicate independent biological replica.

### The C-terminal disulfide bond in OpcA is essential for activation of G6PDH

The interaction between OpcA and monomer C of G6PDH occurs through a loop that is supported by a disulfide bridge between residues C410 and C416 of OpcA (Figure 3D). Two other cysteine residues of OpcA that possibly form another disulfide bond, C192 and C204, are located in the N-terminal domain of OpcA, which is not part of the G6PDH interface (Figure 3A-C). To determine the impact of these covalent interactions on the activation of G6PDH by OpcA, we generated recombinant OpcA variants with mutations C192S and C410S. Analysis of G6PDH activity confirmed that the interface resulting from formation of the C410-C416 bridge is necessary for activation of G6PDH (Figure 6 A). The activity of G6PDH in the presence of the OpcA C410S mutant was comparable to the activity observed in the absence of OpcA. In contrast, the OpcA C192S mutant only partially activated G6PDH (approximately 48 % compared to the WT). Furthermore, in the presence of the C410S/C192S double mutant, which mimics a fully reduced OpcA, the substrate affinity of G6PDH decreased, with a Km of 16.75 mM vs the Km of 8.09 mM determined in the absence of OpcA (Figure 6 B). A similar impact on G6PDH activity occurred when OpcA was pre-incubated with reduced TrxA before the enzymatic assay (Figure 1G), suggesting that reduced OpcA can bind G6PDH and inhibit its activity.

**Fig. 6.**
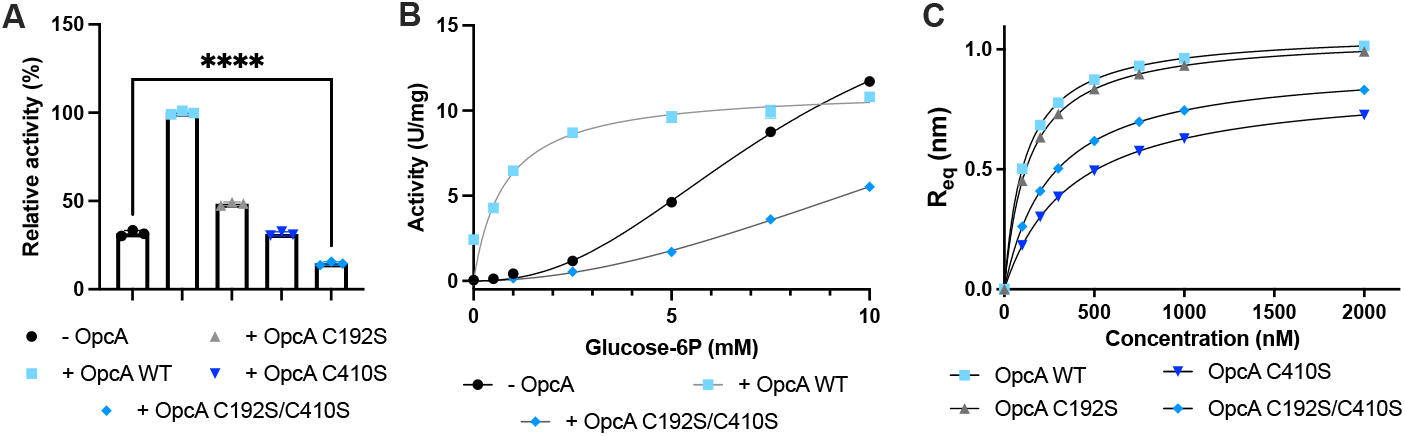
Biochemical characterization of OpcA mutants. (A) Activity of G6PDH in the presence of OpcA mutants. (B) Enzyme kinetics of G6PDH in the absence of OpcA and in the presence of OpcA WT and C192S/C410S mutant. (C) Binding kinetics of OpcA WT and mutants to G6PDH. Every condition was measured at least three times. Error bars represent the SD. Error bars are not visible only when they are smaller than the icon size. Four asterisks represent a p-value ≤ 0.0001.

To confirm this hypothesis, we examined the binding capacity of WT and mutant C410S/C192S OpcA to G6PDH via biolayer interferometry (Figure 6 C). These analyses showed that, although the mutant OpcA variants presented weaker binding to G6PDH than the WT OpcA, they still showed a strong affinity for the enzyme.

### The interaction between G6PDH and OpcA occurs in vivo under different environmental conditions in Synechocystis

Based on the aforementioned findings, OpcA is capable of binding to G6PDH in both its oxidized and reduced state, indicating that this interaction occurs during both heterotrophic phases and photoautotrophic growth within the cell. To further verify this *in vivo*, we used the split NanoLuc technology (NanoBit) to measure the G6PDH-OpcA interaction in changing conditions. This method utilizes a structural complementation reporter system based on a small but highly active luciferase called Nanoluc, which is split into a Large Bit (LgBit) subunit and a small complementary peptide or Small Bit (SmBit). The LgBiT and SmBiT subunits are genetically fused to proteins of interest and expressed in cells. Upon protein interaction, the subunits combine to form an active enzyme, thereby producing a bright luminescent signal in the presence of substrate. This complementation is reversible, which allows the observation of the formation and dissipation of protein-protein interactions (16). We created two *Synechocystis* double mutant strains with *zwf* (the gene encoding for G6PGH) and *opcA* C-terminal tagged with either the LgBit or the SmBit on a genomic level. This way, both genes of interest were expressed under their native promoter (Figure S6). As a negative control, we used a plasmid-encoded fluorescence protein (mVenus) under a strong, rhamnose-inducible promoter tagged C-terminally with SmBit to the *zwf* -LgBit and *opcA*-LgBit mutant, respectively.

We previously demonstrated that cells recovering from nitrogen starvation undergo a heterotrophic phase for about 16 h after the addition of nitrogen, even in the presence of light. They then transition to a mixotrophic phase and eventually resume photoautotrophic growth (17). How these transitions affect the intracellular redox state was so far unclear, but given the gradual switch from respiration to photosynthesis during this process, we expected to see differences in the redox state of OpcA. Similarly, OpcA was expected to be in a reduced state during the day and oxidized during the night. We analyzed the redox state of OpcA in extracts from cells collected during the light and dark phase, and during nitrogen starvation and resuscitation (Figure 7A). During vegetative growth in the light, OpcA was present in different redox states (“light” and “veg”), with most of it being reduced, while in the dark OpcA was mainly oxidized (“dark”). After cells were transferred to nitrogen-free medium, OpcA started to shift towards the oxidized state (“3 d -N”) to reach full oxidation after one month of nitrogen starvation (“1 m - N”). When nitrogen was added to these starved cells, OpcA was progressively reduced (“8 h + N” and “24 h + N”).

**Fig. 7.**
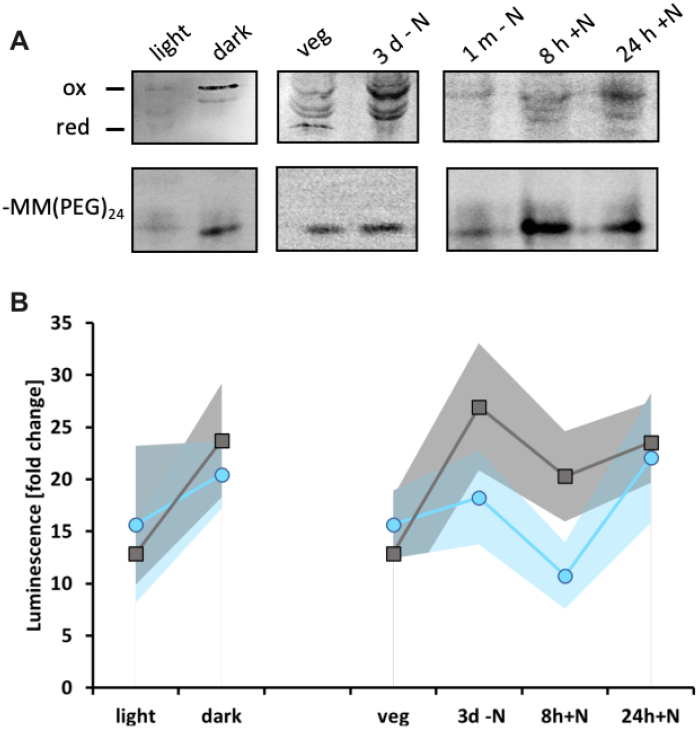
Analysis of the interaction of G6PDH and OpcA *in vivo*. (A) PEGylation assay showing the redox state of OpcA during illuminated, dark, vegetative growth (veg), nitrogen-starved (3d -N, 1 month -N), and resustaction conditions (8h +N, 24h +N). On top, cysteine residues that were involved in a disulfide bridge at the moment of harvest were labeled with MM(PEG)_24_. On the bottom, the same cell extracts without MM(PEG)_24_ labelling are shown. (B) NanoBit assay with G6PDH and opcA C-terminally tagged with either the large (LgBit) or the small (SmBit) fragment of a split NanoLuc luciferase under different conditions (similar to A).

We then measured the interaction between G6PDH and OpcA *in vivo* by recording the luminescence emitted by the NanoBit reporter strains. A significant fold change in luminescence was detected for the OpcA-LgBit/G6PDH-SmBit and G6PDH-LgBit/OpcA-SmBit strains under illumination, darkness, nitrogen starvation, and resuscitation, compared to the control strain (Figure 7B), indicating that G6PDH and OpcA interact under all these conditions. However, differences in fold change could be observed among the different time points. Although the signal diverged between the two mutants, the overall trend was similar: a higher signal was measured in the dark than in the light. This supports our biolayer interferometry data, which shows that the affinity of OpcA for G6PDH is higher when OpcA is oxidized (Figure 6C).

## Discussion

G6PDH catalyzes the first reaction of the OPP pathway, as well as the glucose-6-phosphate and OPP shunts, and plays a key role in the generation of reducing power and metabolic intermediates (4–6, 14). In humans, activation of G6PDH requires binding of a structural NADP^+^ molecule to an allosteric site, which triggers conformational changes that activate the enzyme (18). In photosynthetic organisms, G6PDH is kept in a reduced, inactivated state by the ferredoxin-thioredoxin system in the light, and is oxidized and activated in the dark (19). In chloroplasts, this regulation is achieved by reversible thiol-disulfide interchange within G6PDH itself (20). In constrast, cyanobacterial G6PDH lacks the cysteine residues involved in thiol exchange (Figure S1A) and relies on the OpcA protein to modulate G6PDH activity. In this work, we found that OpcA carries out the dual function of allosterically stabilizing and activating/inhibiting G6PDH, thereby rendering it susceptible to redox regulation.

Our cryo-EM analysis revealed the mechanism of G6PDH allosteric regulation by OpcA. The G6PDH-OpcA structure shows that the C-terminus of OpcA interacts with the C-terminal beta-alpha domain of G6PDH (Figure 3). Here, the interaction surface on G6PDH monomer C corresponds to the structural NADP^+^ allosteric binding site of the human protein, which is essential for the enzymatic activity (14, 21) (Figure S5C). In cyanobacteria, binding of OpcA causes conformational changes of the G6PDH tetramer, particularly affecting the substrate binding site (Figure 4) and activates G6PDH by increasing its affinity of G6PDH for glucose-6-phosphate (Figure 1B). This is supported by the reorganization of H197 and Y198, the key residues involved in ligand binding and catalysis (Figure 4B-E). In the human enzyme, similar rearrangements of homologous H201 and Y202 have been observed upon binding of the structural NADP^+^ molecule to G6PDH (14), suggesting a comparable mechanism of allosteric regulation. OpcA also interacts with the N-terminus of one of G6PDH monomers, allowing its structural organization (Figures 3C,E and 4B). In the absence of OpcA, this N-terminal fragment is unordered and not resolved in the structure. The function of this segment is so far unknown, but it may play a role in G6PDH regulation or in supporting the interaction with OpcA. In our cryo-EM studies, we detected only G6PDH-OpcA complexes formed by a G6PDH tetramer bound to a single OpcA protein. This may not represent a fully saturated and active state since our further analysis of the enzymatic activity and molecular mass of the complex under different OpcA concentrations suggests that up to four OpcA monomers can bind to the G6PDH tetramer and even further stimulate its activity (Figure 5). However, given that G6PDH seems to be more abundant than OpcA intracellularly, full saturation is probably not reached *in vivo*. This situation might result in a mixed population of G6PDH tetramers that bind different numbers of OpcA molecules and therefore display distinct activation levels. A G6PDH population that is partly active also in the light might be beneficial to minimize futile cycling with the Calvin-Benson cycle while at the same time allowing limited flux via the OPP shunt into the cycle. For example, upon adding glucose in the light, which represents highly reducing conditions, *Synechocystis* feeds about 9 % of its carbon flux via the OPP shunt into the Calvin-Benson cycle, showing that G6PDH has to be at least partly active under these conditions (22).

The interface of OpcA with the allosteric site of G6PDH involves a C-terminal loop containing a disulfide linkage between C410 and C416 (Figure 3C-E). Even though the presence of this disulfide bond under oxidizing conditions does not seem to be required for formation of the OpcA-G6PDH complex (Figure 6C), it has a strong effect on G6PDH activity (Figure 6A,B). The formation of the C410-C416 disulfide bond apparently induces a conformational change in OpcA, which enables allosteric activation of G6PDH through a structural alteration of its substrate binding site (Figure 4). This way, a redox control of the OPP pathway is implemented in cyanobacteria. During photoautotrophic growth, the thioredoxin system maintains the OpcA mostly in a reduced state, which binds and inhibits G6PDH to minimize carbon catabolism via the OPP pathway. Under nitrogen starvation, cells have degraded most of their photosynthetic machinery and electron transport is kept to a minimum (17), which causes OpcA to progressively switch towards an oxidized state making it capable of activating G6PDH. Upon the recurrence of nitrogen availability, cells rely on the OPP pathway to obtain the energy and metabolites needed to restore the photosynthetic machinery (23). During this period, we observed a progressive reduction of OpcA as the cells regained photosynthetic capacity. This analysis shed light on the intracellular redox state of *Synechocystis* cells during these transitions, which is of crucial importance to understand the regulatory mechanisms behind this developmental program.

In conclusion, this study has revealed a distinct mechanism of redox regulation of the OPP pathway and OPP shunt in cyanobacteria, which involves the OpcA protein. Curiously, OpcA is not exclusive to these photoautotrophic microbes, but it is also present in actinobacteria. Interestingly, in higher photosynthetic organisms, which lack OpcA, the redox sensitivity of OpcA has been integrated in G6PDH itself. OpcA binds to G6PDH both in its oxidized and reduced form and either activates or inhibits enzyme activity. As the activity of G6PDH depends on the concentration of OpcA, and as full saturation is not reached *in vivo* according to our data, this mechanism allows a delicate fine-tuning of the enzyme. Furthermore, our functional and structural analyses have shown that the function of OpcA is not limited to redox control, but also includes an allosteric stabilizing-activating effect on G6PDH. In mammals, this role seems to have been overtaken by the regulatory structural NADP^+^ molecule, as it has been shown for the human protein. Future studies will exploit our findings to further investigate this unique redox regulatory mechanism and determine whether OpcA has additional functions and targets beyond G6PDH.

## Materials and Methods

### Protein overexpression and purification

*Escherichia coli* Rosetta-gami (DE3) was used for protein overexpression. All primers and plasmids used are shown in Table S2 and Table S3, respectively. Cells were cultivated in LB medium (1 L of culture in 5 L flasks) at 37 °C until they reached exponential growth (*OD*_600_ 0.6-0.8) and protein overexpression was then induced by adding either 0.5 mM IPTG (for His-tagged proteins) or 75 μg/L anhydrotetracycline (for Strep-tagged proteins), followed by incubation at 20°C for 16 h. Cells were harvested by centrifugation at 4000 g for 10 min at 4 °C, and disrupted by sonication in 40 mL of lysis buffer (100 mM Tris-HCl pH 7.5, 150 mM KCl, 5 mM MgCl_2_, 10 mM imidazole (only for His-tagged proteins), DNAse, and protease inhibitor cocktail (COmplete, Sigma-Aldrich, Taufkirchen, Germany). The cell lysates were centrifuged at 20,000 g for 1 h at 4°C and the supernatants were filtered with a 0.22 μm filter. For the purification of His-tagged proteins, 5 mL Ni-NTA HisTrap columns (GE Healthcare, Chicago, IL, USA) were used. The cell extracts were loaded into the columns, washed with wash buffer (100 mM Tris-HCl pH 7.5, 150 mM KCl, and 50 mM Imidazole) and eluted with elution buffer (100 mM Tris-HCl pH 7.5, 150 mM KCl, and 500 mM Imidazole). For the purification of Strep-tagged proteins, 5 mL Ni-NTA Strep-tactin superflow (Qiagen, Germantown, MD, USA) columns were used. The cell extracts were loaded into the columns, washed with wash buffer (100 mM Tris-HCl pH 7.5 and 150 mM KCl) and eluted with elution buffer (100 mM Tris-HCl pH 7.5, 150 mm KCl, and 2.5 mM desthiobiotin). The buffer of all purified proteins was exchanged via dialysis to 100 mM Tris-HCl pH 7.5, 150 mm KCl, and 5 mM MgCl_2_ in a 3.5 kDa cutoff dialysis tube (Spectrum Laboratories, Los Angeles, CA, USA). All purifications were checked via SDS-PAGE.

### Measurement of G6PDH activity in vitro

The reaction buffer was composed of 100 mM Tris-HCl pH 7.5, 150 mM KCl, 10 mM MgCl_2_ and 1 mM NADP^+^. 500 ng G6PDH were added to each reaction. When indicated, OpcA was added at the described molar ratio. When stated, the enzymes were pre-treated with DTT_*red*_, DTT_*ox*_ or CuCl_2_ at the concentration indicated in the figure legends for 30 min. The reaction was started by the addition of glucose-6-phosphate at the indicated concentrations. Absorption change at 340 nm was continuously measured for 15 min at 30 °C. The enzymatic activity was then calculated. At least three replicates were measured for each condition.

### Size exclusion chromatography multiangle light scattering (SEC– MALS)

SEC-MALS experiments were performed using and Ä KTA purifier system connected to a Superose 6 Increase 10/300 GL column (GE healthcare) at a flow rate of 0.4 ml/min in running buffer (100 mM Tris-HCl pH 7.5, 150 mM KaCl, and 10 mM MgCl_2_). The column was calibrated using the gel filtration calibration kit LMW and HMW (GE Healthcare) according to the manufacturer’s instructions. To analyze the oligomeric state of the recombinant proteins, the ÄKTA system was connected to a downstream multi angle light scattering (MALS) system using the miniDAWN TREOS combined with an Optilab T-rEX refractometer (Wyatt Technology, Dernbach, Germany). Data analysis was performed using the software ASTRA 7 (Wyatt Technology) and Unicorn 5.20 (Build 500) (General Electric Company, Boston, MA USA).

### Biolayer interferometry using the Octet K2 system

Protein-protein interaction was tested *in vitro* by biolayer interferometry using the Octet K2 system (Satorius, Göttingen, Germany). All experiments were performed in HEPES buffer (100 mM HEPES-KOH pH 7.5 and 10 mM MgCl_2_). His8-tagged G6PDH was immobilized on Ni-NTA sensor tips (Satorius) by exposing the sensors to a 500 nM solution of G6PDH for 120 s (loading), followed by a 60 s baseline measurement. To avoid unspecific binding, the sensor tips were then dipped in a solution containing 600 nM of His8-PII-ΔTloop, followed by a second 60 s baseline measurement. For the binding of OpcA, the sensor tips were dipped in solutions with different concentrations of Strep-OpcA WT or mutants, as indicated (association). The assay was finalized with a 120 s dissociation step. As a control, the loading was done using His8-PII instead of His8-G6PDH. The biosensors were regenerated after each use with 10 mM glycine (pH 1.7) and 10 mM NiCl_2_ as proposed by the manufacturers. The recorded curves were aligned to the baseline before the association step.

### Mass photometry analysis

Microscope coverslips (No. 1.5H, 24×5, Marienfeld, Germany) were cleaned by 3 cycles of sequential immersion in isopropanol (HPLC grade) and Milli-Q water. CultureWell gaskets (3 mm DIA x 1 mm Depth, Grace Bio-Labs, Bend, OR, USA) were assembled on the clean coverslips. Samples were diluted with buffer (50 mM Tris-HCl pH 7.5, 5 mM MgCl_2_) to working concentrations of 100-150 nM. Acquisition was done in a TwoMP mass photometer (Refeyn, Oxford, UK) using the AcquireMP v2022 R1 software. A calibration measurement was performed before each experiment. To find focus, 5 μL of buffer were added to a gasket’s well, and the focal position was identified and secured in place with an autofocus system based on total internal reflection for the entire measurement. For acquisition, 5 μL of diluted protein (standard or sample) was added to the same well, autofocus was stabilized, and movies of 60 s duration were recorded. Each sample was measured at least three times independently. Images were processed and analyzed using the DiscoverMP v2022 R1 software as described in Kofinova et al. (2024)(24).

### PEGylation assay

*Synechocystis* cultures (50 mL) grown under different conditions were supplemented with 10 mM N-ethyl-maleimide (NEM, Sigma-Aldrich) and further cultivated for 1 h to block free cysteine residues. Cells were then collected by centrifugation (4,000 g for 10 min at 4 °C). Cell pellets were resuspended in 300 μL of buffer A (50 mM Tris-HCl pH 7.5, 50 mM NaCl, 100 mM NEM) and 50 μL of glass beads (0.25 - 0.5 mm, Roth, Karlsruhe, Germany) were added. Cells were lysed using a MP FastPrep-24 5G homogenizer (MP Biomedicals, Irvine, CA, USA). Samples were then centrifuged at 15000 g for 15 min at 4 °C and supernatants were transferred to fresh tubes. Proteins were precipitated by adding 10 % TCA, incubating for 1 h on ice, centrifuging at 15000 g for 15 min and washed with acetone. Protein pellets were resuspended in buffer B (50 mM Tris-HCl pH 7.5, 50 mM NaCl, 2 % SDS, 100 mM DTT) and incubated for 1 h on ice to reduce disulfide bonds. Protein precipitation was repeated and pellets were then resuspended in 50 μL buffer C (50 mM Tris-HCl pH 7.5, 50 mM NaCl, 2 % SDS, 7.5 % glycerol, 0.01 % bromophenol blue) and divided into two 25 μL samples. 10 mM MM(PEG)_24_ was added to one of the samples to label cysteines, the other sample was used as a control. Proteins were separated by polyacrylamide gel electrophoresis (PAGE) and transferred to a BioTrace™ polyvinylidene fluoride membrane (Pall Corporation, New York, NY, USA) using a semi-dry blotting system (PEQLAB, Wilmington, CA, USA) at 20 V for 30 min. Membranes were blocked in TBS-T buffer (50 mM Tris, 150 mM NaCl, 0.1 % Tween-20, pH 7.5) with 5 % milk powder for 1 h at room temperature, washed with TBS-T buffer and incubated with mouse-produced monoclonal anti-polyHistidine-Peroxidase antibodies (Sigma-Aldrich) in a 1:1000 dilution in TBS-T buffer overnight at 4 °C. Membranes were washed again and visualized by Lumi-Light Plus detection reagent (Roche, Basel, Switzerland).

### Statistical analysis

Statistical details for each experiment can be found in the figure legends. GraphPad PRISM was used to perform paired Student’s t-tests to determine the statistical significance. Asterisks in the figures were used to symbolize the p-value: One asterisk represents p ≤ 0.05, two asterisks p ≤ 0.01, three asterisks p ≤ 0.001, and four asterisks p ≤ 0.0001.

### Synechocystis sp. PCC 6803 mutant creation

To generate strains in which OpcA and G6PDH are tagged with either one of the subunits of the NanoBit system (large bit: LgBit and small bit: SmBit (16) or a 6xHis-Tag, so-called target plasmids were synthesized (GENEWIZ, Leipzig, Germany) that contained the following elements: 250 bp upstream of the C-terminal end of the gene of interest without the STOP codon, BamHI and EcoRI restriction sites, and 250 bp downstream of the site of insertion in a pUC57-Simple backbone. A separate plasmid containing a linker region with a TEV cleavage site and a 6x-His tag (pTK0XX-Fed9-His (25)) was cut with NheI to insert the LgBit via Gibson assembly resulting in the plasmid pTK0XX-Fed9-lBit. The SmBit, including the described linker region (16) was synthesized and ligated into pTK01-Fed9-His cut with EcoRI and EcoRV, resulting in pTK30-Fed9-sBit. These plasmids were then cut with EcoRV to add different antibiotic-resistance cassettes. The fragments containing the various tags and antibiotic resistance cassettes were then cut out with BamHI and EcoRI and ligated into the target plasmids, which were also cut with BamHI and EcoRI. Finally, these plasmids were transformed into *Synechocystis* to create single and double mutants containing both tagged genes. The negative control stains for the NanoBit assay were created by adding the same linker region and SmBit used above to an RSF1010-based plasmid containing a rhamnose inducible promotor (26) cut with NdeI and StuI. Then, the GFP-based fluoresce protein mVenus was amplified and added in front of the SmBit tag in a Gibson assembly. This plasmid was finally transformed via electroporation into the single mutants opcA-LgBit and ZWF-G6PDH-LgBit. A list of all primers and plasmids used in this study is provided in Table S2 and Table S3.

### NanoBit assay

Double mutants and negative controls were pre-cultivated in 50 ml BG_11_ in shaking flasks in constant light (50 μmol photons / *m*^2^*s*^2^) at 28 °C for several days. Then, each culture was diluted to an OD_750_ of 0.1, transferred to three test tubes, and further cultivated until the OD_750_ was above 1. One mL of each sample was taken, centrifuged (4000g, 5 min) and resuspended in BG_11_ to an OD_750_ of 1. The remaining cells were centrifuged (4000 g, 5 min) and resuspended in BG_11*−*0_ (BG_11_ without NaNO_3_) three times to create nitrogen starvation conditions. After three days, samples were taken as described before. The remaining cells were then centrifuged and resuspended in BG_11_ and further cultivated. Samples were also taken after 8 h and 24 h during resuscitation from nitrogen starvation. All samples were analyzed shortly after being taken and were kept in the light to ensure sufficient oxygenation. 100 μL of cells of each culture were added to a white flat bottom 96-well plate. Then, 25 μL of diluted Nano-Glo reagent (Promega, Walldorf, Germany) was added. Cells were illuminated for at least 5 min (50 μMol photons / *m*^2^*s*^2^) before measuring the luminescence every 80 s for 15 min in darkness using a CLARIOstar plate reader (BMG-Labtech, Ortenberg, Germany).

### Protein extract preparation, polyacrylamide gel electrophoresis, and immunoblotting

Cells were harvested by centrifuging and resuspended in ACA buffer (750 mM -amino caproic acid, 50 mM BisTris/HCl, pH 7.0, 0.5 mM EDTA). Cells were broken by vortexing with glass beads (0.17 – 0.18 μm diameter) for 2 min at full speed at 4 °C. Glass beads and cell debris were pelleted by centrifuging at 3,500 xg for 10 min. Chlorophyll concentrations were determined by extraction with methanol and measuring the absorption at 656nm of. In SDS PAGE immunoblotting analysis, an amount corresponding from 0,25 μg to 0,025 μg of Chl were loaded per lane. These samples were solubilized at RT for 1h in 1x Laemmli sample buffer and separated on 12.5 % (w/v) polyacrylamide BisTris gels using a MES running buffer (250 mM MES, 250 mM Tris, 5 mM EDTA, 0.5 % (w/v) SDS). The resultant gels were electroblotted onto nitrocellulose membrane. For immunoblotting 5 % (w/v) milk powder in 1x PBS-T was used as a blocking solution, and all washing steps were performed with 1xPBS-T or PBS. The immunoblotting analyses were conducted using mouse anti-6xHis-primary antibodies (Genescript, New Jersey, USA) and horseradish peroxidase-conjugated secondary antibodies Peroxidase-conjugated Anti-mouse (Jackson ImmunoResearch Europe Ltd., Cambridge, UK). After detecting the luminescent signal using a CCD-camera-equipped imager system (Azure 400, Azure biosystems, Dublin, USA), all transferred protein was stained by ponceau staining and also detected with the imager system. All images were analyzed using AzureSpotPro software (Azure 400, Azure biosystems, Dublin, USA).

### Negative-stain EM analysis

Negative-stain electron microscopy was employed to analyze the sample of purified G6PDH-OpcA. 3 μl of the sample at a protein concentration of around 0.06 mg/ml was deposited onto freshly glow-discharged carbon-coated copper grids with plastic support. The grids were then blotted and stained using a 2 % (w/v) uranyl formate solution, following the standard protocol (27). The micrographs (Figure S2A) were manually collected using a JEM-2100Plus transmission electron microscope (JEOL) operating at 200 kV, equipped with a XAROSA CMOS 20-megapixel camera (EMSIS), at a nominal magnification of 30,000 (3.12 Å per pixel). Subsequent data analysis was done using ImageJ (28) and cryoSPARC (10).

### Cryo-EM sample preparation and data acquisition

For cryogenic electron microscopy (cryo-EM), 3 μl of the purified G6PDH-OpcA sample at a protein concentration of approximately 1 mg/ml were applied to glow-discharged C-Flat grids (R1.2/1.3 3Cu-50) (EMS) and directly plunge-frozen in liquid ethane using a Vitrobot Mark IV (Thermo Fisher Scientific) with the environmental chamber set at 100 % humidity and 4°C. Movies (6483) were automatically acquired using EPU (Thermo Fisher Scientific) on a Glacios cryogenic transmission electron microscope (Thermo Fisher Scientific) operating at 200 kV with a Selectris energy filter and a Falcon 4 detector (both Thermo Fisher Scientific). Data recording was performed in Electron Event Representation (EER) mode at a nominal magnification of 130,000 (0.924 Å per pixel) in the defocus range of -0.8 to -1.8 μm, with an exposure time of 5.65 s, resulting in a total electron dose of approximately 50 e-Å ^*−*2^.

### Cryo-EM image processing

All cryo-EM data underwent preprocessing in cryoSPARC Live, followed by additional processing in cryoSPARC v4 (10) (see Figures S2 and S3 for details). Preprocessing in cryoSPARC Live included motion correction, contrast transfer function (CTF) estimation, micrograph curation, and particle selection using a blob picker, followed by 2D classification. Further steps performed in cryoSPARC involved particle picking using templates derived from cryoSPARC Live preprocessing, 2D classification, duplicate particle removal, and ab-initio reconstruction followed by heterogeneous refinement using 9 classes (Figure S3). The two best classes were further processed separately. One class (435944 particles), corresponding to the complex of G6PDH-OpcA, underwent a round of NU-refinement (29). This step was followed by 3D classification with three classes. The best 3D class was then subjected to a round of

NU-refinement followed by a local refinement using a mask covering the entire structure. This resulted in a G6PDH-OpcA map of 3.70 Å (Gold Standard Fourier Shell Correlation (GSFSC) value of 0.143). Another best class (373874 particles) from the first heterogeneous refinement with nine classes corresponded to an apo state of G6PDH. This class underwent NU-refinement (29) followed by ab-initio reconstruction and heterogeneous refinement with three classes. The two best classes from this heterogeneous refinement (210008 and 148324 particles), combined with the best class from another heterogeneous refinement run with particles (435944) originating from the class initially assigned to the G6PDH-OpcA class, were subjected to another round of heterogeneous refinement with three classes. The best class from this heterogeneous refinement was further used for NU-refinement with C1 symmetry, followed by NU-refinement with C2 symmetry and NU-refinement with C2 symmetry and defocus refinement. This resulted in a G6PDH map of 3.27 Å (Gold Standard Fourier Shell Correlation (GSFSC) value of 0.143). Unsupervised B-factor sharpening was applied and local resolution estimation was performed within cryoSPARC (Figure S2).

### Model building and refinement

AlphaFold structure predictions of *Synechocystis* G6PDH and OpcA (Uniprot: P73411, P73720, respectively) were manually fitted into the consensus and local-refinement maps. The models of G6PDG and OpcA underwent manual adjustments and refinements in Coot (30). Iterative rounds of real-space refinement in PHENIX (31), accompanied by manual adjustments in Coot, were performed. Model validation was carried out using MolProbity (32) in PHENIX (see Table S1 for model refinement and validation statistics). Visualization and figure preparation were performed in UCSF ChimeraX (33), and Affinity Designer 2.

### Data availability

The cryo-EM density maps and associated atomic models presented in this work have been submitted to the Electron Microscopy Data Bank and Protein Data Bank, where they can be accessed using the numbers EMD-xxx and PDB-xxx, respectively.

## Acknowledgments

We thank Eva Nußbaum and Heinz Grenzendorf for their assistance with cyanobacterial cultivation and protein purification. We acknowledge Prof. Dr. Arne Moeller for access to the EM platform at the CellNanOs center at Osnabrü ck University and support in cryo-EM data analysis. We also thank Dr. Dovile Januliene and Kilian Schnelle for their advice, Dr. John Weir and Maria Kharlamova for their help with mass photometry experiments, and Dr. Libera Lo Presti for her assistance editing this manuscript. This work was supported by the infrastructural funding via the Cluster of Excellence (EXC2124) “Controlling Microbes to Fight Infections” at the University Tübingen, and the German Research Council (DFG) FOR 2816, SFB 1557, DFG INST190/196-1 FUGG, MO2752/3-6, and GU1522/5-1.

