## Supplementary Information for "Structural basis of the allosteric regulation of cyanobacterial glucose-6-phosphate dehydrogenase by the redox sensor OpcA"

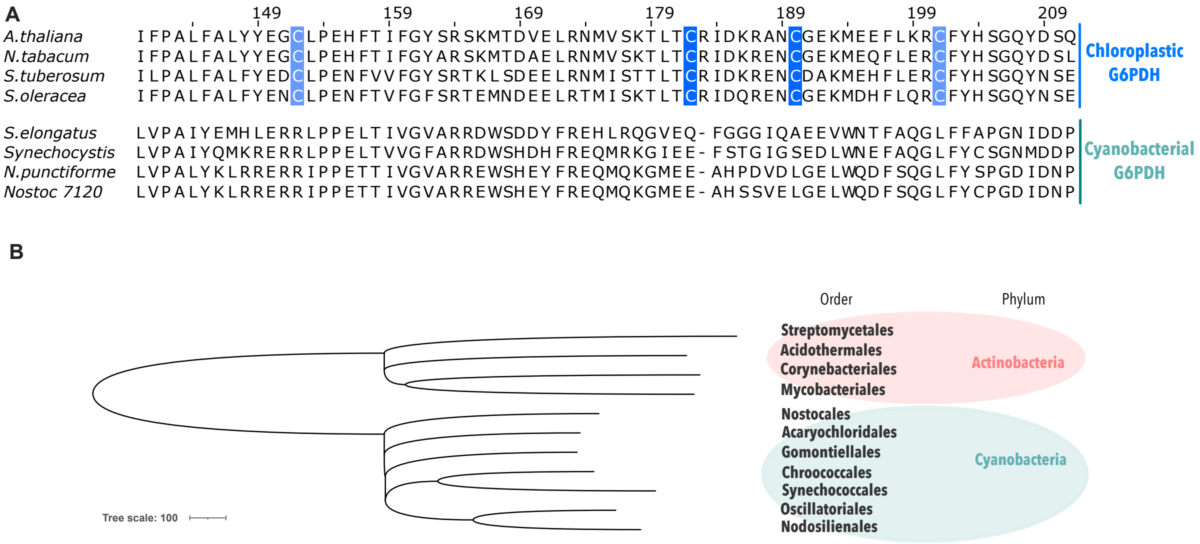


**Figure S1.** Conservation of G6PDH and OpcA. (A) Alignment of different chloroplastic and cyanobacterial G6PDH. The redox reactive cysteine residues are highlighted in blue. (B) Phylogenetic tree of selected species from the phyla Cyanobacteria and Actinobacteria. Sequences were obtained from the UniProt database. Alignment and phylogenetic tree data were obtained using NCBI Constraint-based Multiple Alignment Tool (COBALT) and edited with Jalview and the Interactive Tree Of Life (iTOL) v5 online tool.

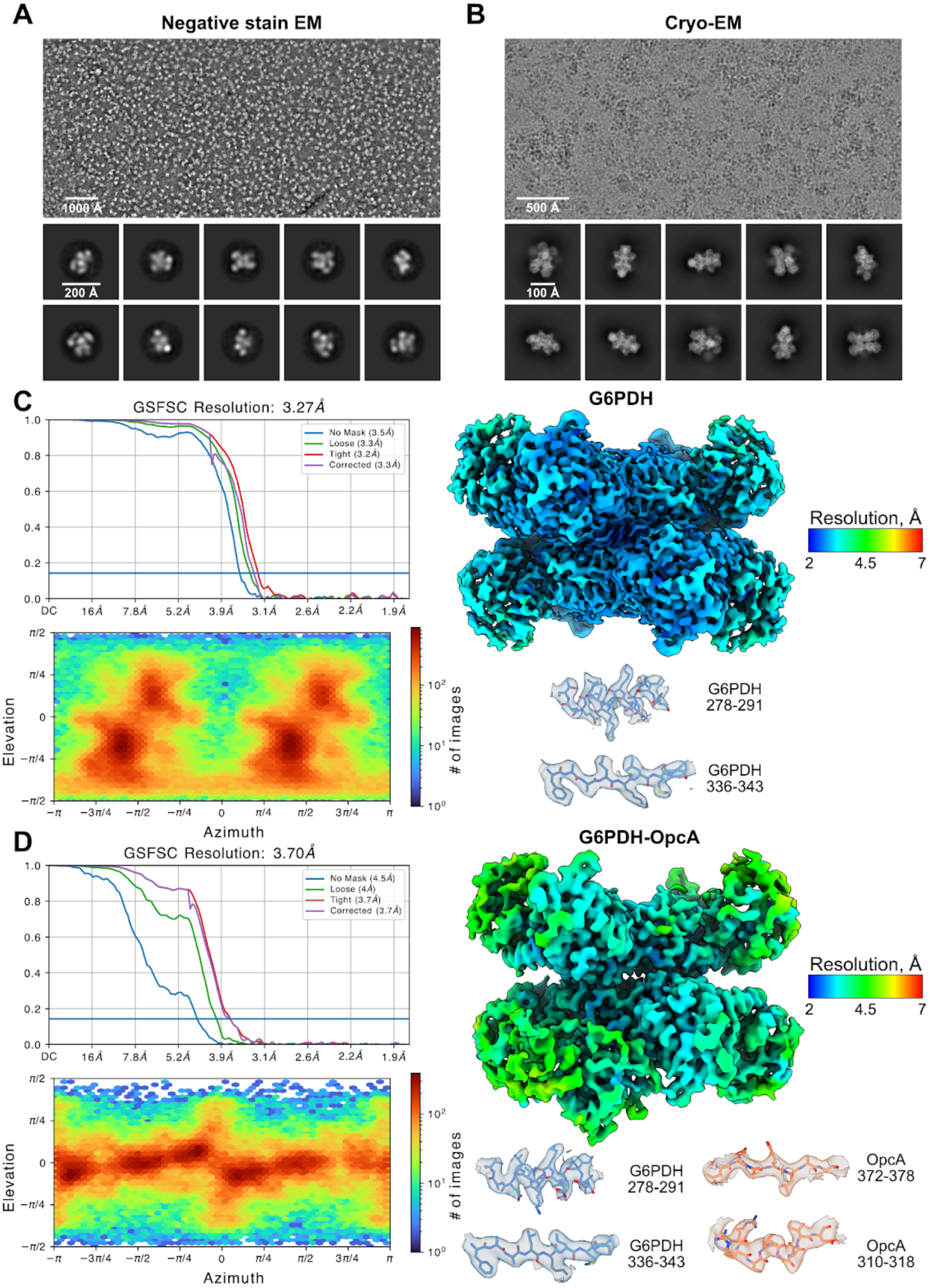


**Figure S2**. Electron microscopy single-particle analysis and structure validation. (A, B) Representative negative stain EM (A) and cryo-EM (B) micrographs and 2D class averages (bottom). (C, D) Gold standard Fourier shell correlation (GSFSC) curve, angular distribution heatmap plot, local resolution estimation generated in cryoSPARC, and model/map fit of selected areas for the final map of G6PDH (C) and G6PDH-OpcA complex (D).

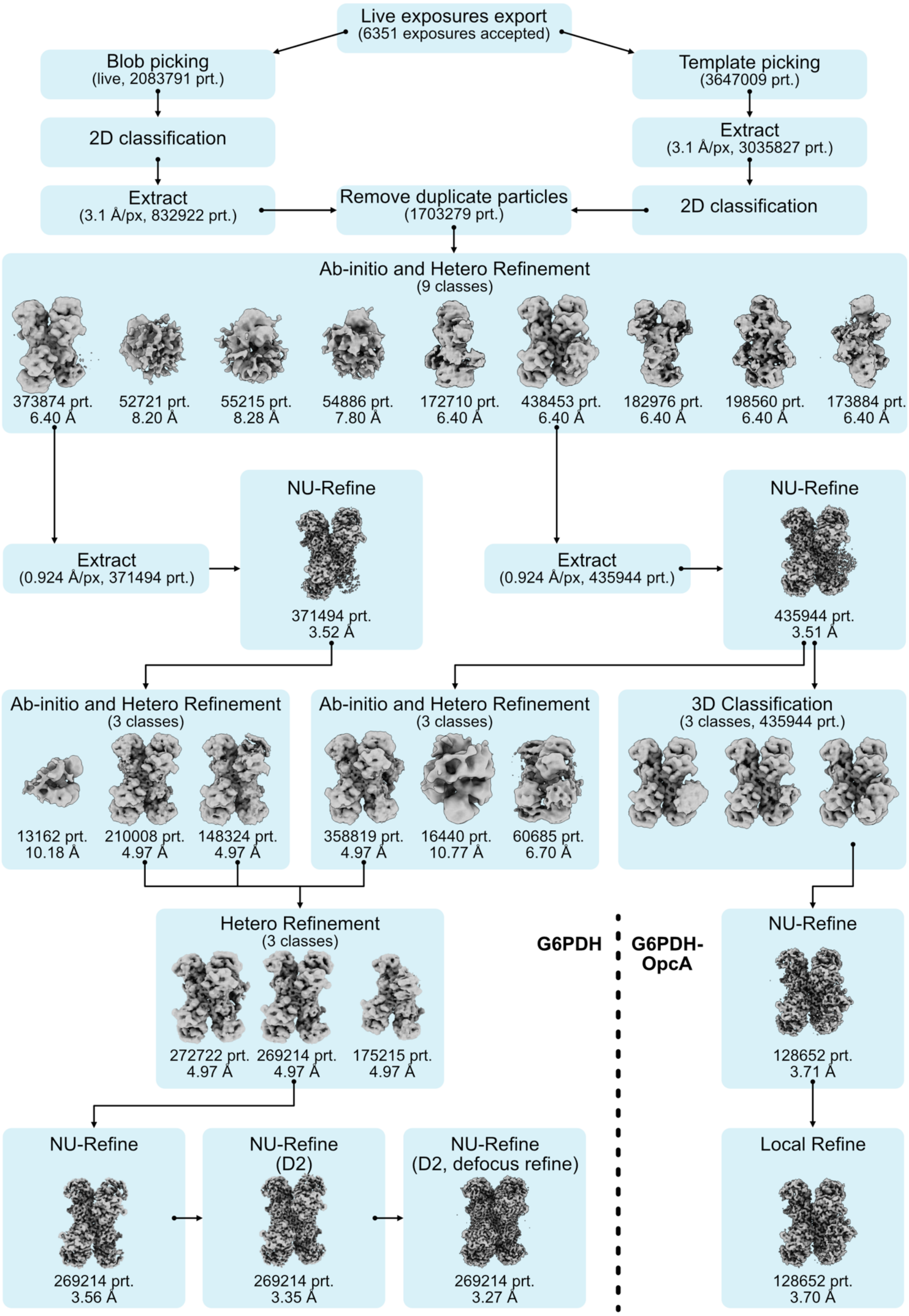


**Figure S3**. Cryo-EM data processing scheme of G6PDH and G6PDH-OpcA. The main steps of the data processing pipeline in cryoSPARC are shown. For Extract jobs, the numbers of particles used and the pixel size are indicated. For maps in each Refinement job, the number of particles, and the achieved resolution are provided.

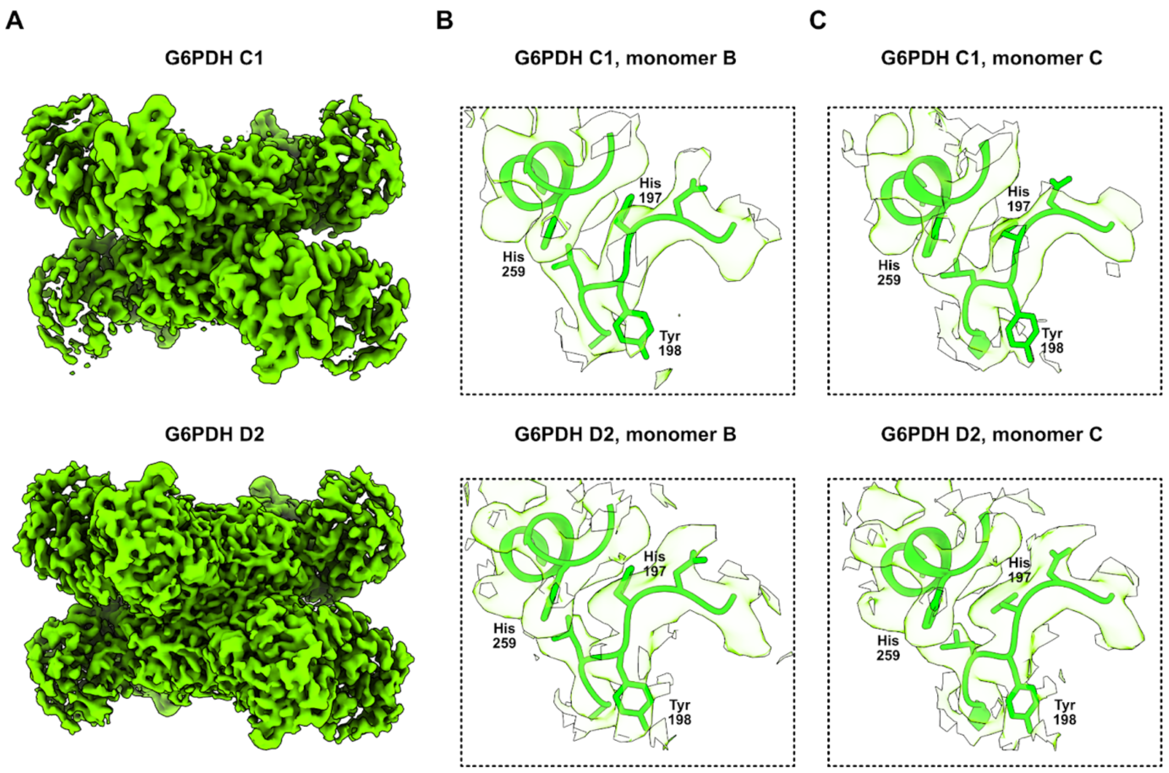


**Figure S4**. Cryo-EM map of apo G6PDH processed without symmetry and with D2 symmetry applied. (A) Comparison of full cryo-EM maps without (top) and with D2 symmetry (bottom). (B,C) Segments of monomer B (B) and monomer C (C) in the region of the enzyme active site (see Figure 4) with the nonsymmetric (upper panels) and D2-symmetric (bottom panels) cryo-EM map zoned around the model.

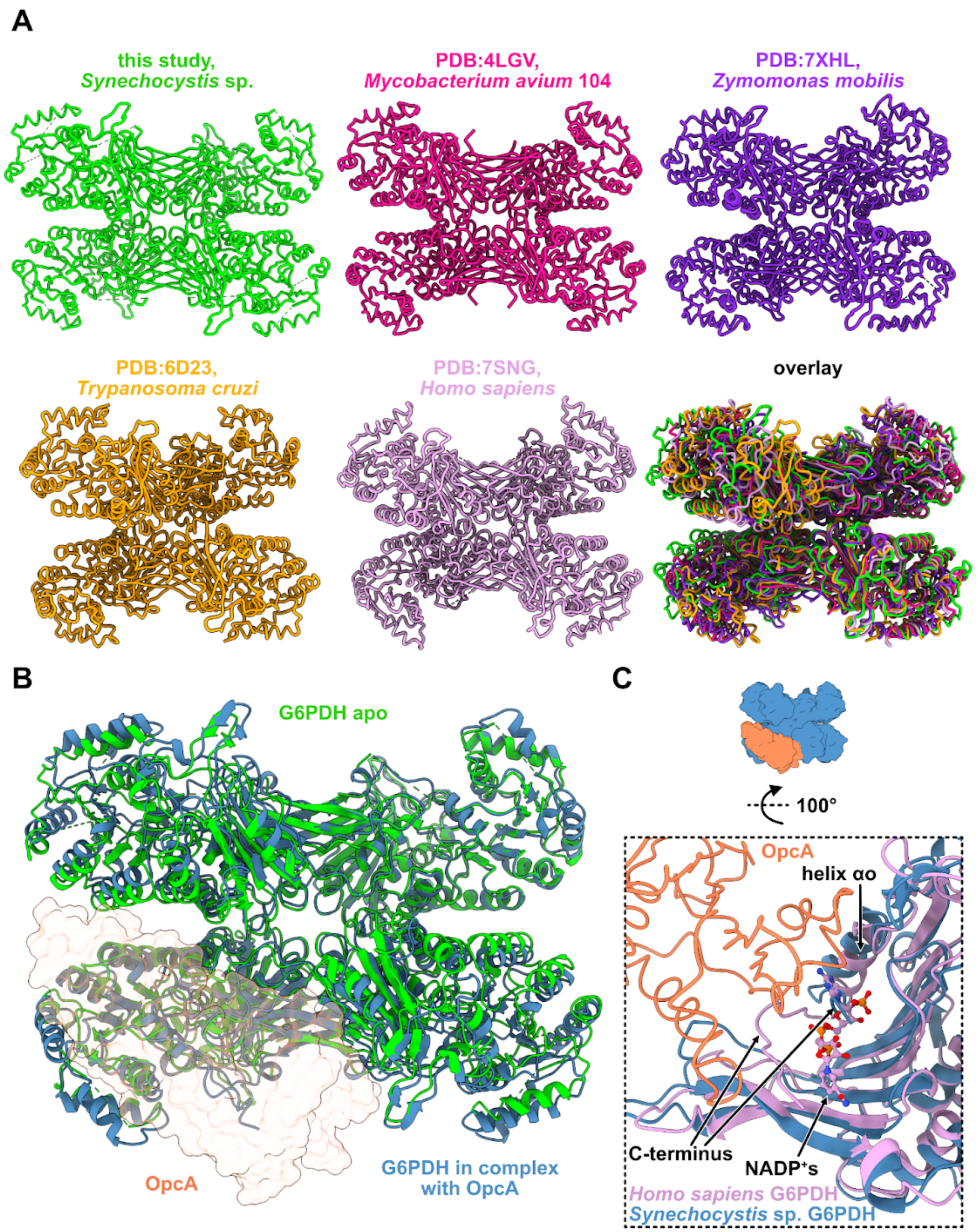


**Figure S5**. Comparative structural analysis of G6PDH. (A) Structural comparison of the *Synechocystis* G6PDH apo tetramer with homologous proteins from different species (the PDB accession codes and organism names are provided). (B) Superposition of the *Synechocystis* G6PDH apo tetramer (green) with the G6PDH (dark blue) from the complex with OpcA (shown as orange semi-transparent surface). (C) Close-up view comparing the allosteric structural $NADP^{+}$s binding site in human G6PDH (PDB: 1QKI; colored plum) with *Synechocystis* G6PDH (colored dark blue) at the binding interface with OpcA (colored orange). Structural $NADP^{+}$ molecule bound to human G6PDH is depicted as balls and sticks.

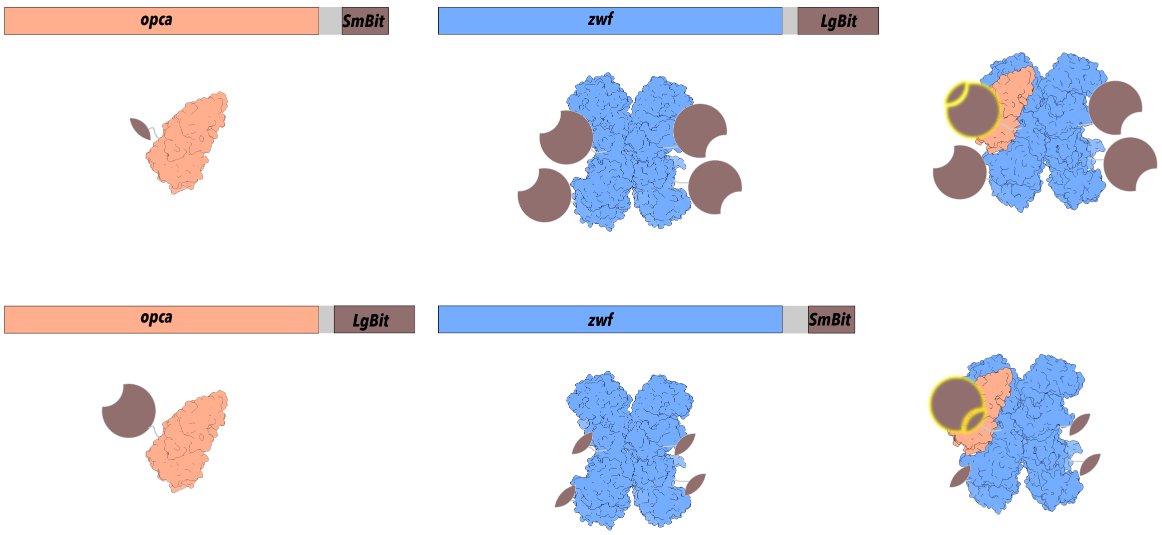


**Figure S6**. Scheme of the constructs created using the NanoBit technology.

**Table S1.** Cryo-EM data collection, refinement and validation statistics.

| | G6PDH C1 (EMDB-xxxx) | G6PDH D2 (EMDB-xxxx) (PDB xxxx) | G6PDH-OpcA (EMDB-xxxx) (PDB xxxx) |
|---|---|---|---|
| **Data collection and processing** | | | |
| Magnification | | 130,000 | |
| Voltage (kV) | | 200 | |
| Electron exposure (e–/Å$^2$) | | 50 | |
| Defocus range (μm) | | -0.8 to -1.8 | |
| Pixel size (Å) | | 0.924 | |
| Symmetry imposed | C1 | D2 | C1 |
| Initial particle images (no.) | | 1703279 (after duplicates removal) | |
| Final particle images (no.) | 269214 | 269214 | 128652 |
| Map resolution (Å) | 3.4 | 3.3 | 3.7 |
| FSC threshold | 0.143 | 0.143 | 0.143 |
| **Refinement** | | | |
| Initial model used | | AlphaFold Uniprot: P73411 | AlphaFold Uniprot: P73411, P73720 |
| Model resolution range (Å) | | 3.3 | 3.7 |
| FSC threshold | | 0.143 | 0.143 |
| Map sharpening *B* factor (Å$^2$) | -168.7 | -137.9 | -111.3 |
| Model composition | | | |
| Non-hydrogen atoms | | 14704 | 18401 |
| Protein residues | | 1833 | 2305 |
| R.m.s. deviations | | | |
| Bond lengths (Å) | | 0.006 | 0.005 |
| Bond angles (°) | | 0.802 | 0.707 |
| Validation | | | |
| MolProbity score | | 1.89 | 2.10 |
| Clashscore | | 6.69 | 9.17 |
| Poor rotamers (%) | | 0.51 | 0.46 |
| Ramachandran plot | | | |
| Favored (%) | | 91.02 | 87.41 |
| Outliers (%) | | 0.78 | 0.66 |

| Table S2. List of used primers | |
|---|---|
| Primer | Sequence (5' - 3') |
| G6PDH fw ol pET15b | CAGCAGCGGCCTGGTGCCGCGCGGCAGCCATATGCTCGAGATGGTAACGCTACTCGAAAATC |
| G6PDH rv ol pET15b | CCAACTCAGCTTCCTTTCGGGCTTTGTTAGCAGCCGGATCCTAAAGTCGGCGCCAGCGACG |
| OpcA fw ol pET15b | CAGCAGCGGCCTGGTGCCGCGCGGCAGCCATATGCTCGAGATGGGGGGAAAGTATCAGCGT |
| OpcA rev ol pET15b | CCAACTCAGCTTCCTTTCGGGCTTTGTTAGCAGCCGGATCTTATCCTGCCTGGGATAACTG |
| G6PDH fw ol pASK | AAATGGCTAGCTGGAGCCACCCGCAGTTCGAAAAAGGCATGGTAACGCTACTCGAAAATC |
| G6PDH rev ol pASK | CGGGTACCGAGCTCGAATTCGGGACCGCGGTCTCGGCCTAAAGTCGGCGCCAGCGAC |
| OpcA fw ol pASK | ACAAATGGCTAGCTGGAGCCACCCGCAGTTCGAAAAAGGCATGGGGGGAAAGTATCAG |
| OpcA rev ol pASK | CCCGGGTACCGAGCTCGAATTCGGGACCGCGGTCTCGGCTTATCCTGCCTGGGATAACTG |
| OpcA-C410S fw | ACCGTACTGTcCTCCGGCACTG |
| OpcA-C410S rev | GCCGCATTGGGCTTGGGT |
| OpcA-C191S fw | TCCGCCTACTcCCCGATCCAAAAG |
| OpcA-C191S rev | AAGCTGGGCCTGTACCCC |

| Table S3. List of used plasmids | |
|---|---|
| Plasmid | Purpose |
| pET15b-G6PDH | Expression of N-terminus His-tagged *Synechocystis* G6PDH in *E. coli* |
| pET15b-OpcA | Expression of N-terminus His-tagged *Synechocystis* OpcA in *E. coli* |
| pASK IBA5-G6PDH | Expression of N- terminus Strep-tagged *Synechocystis* G6PDH in *E. coli* |
| pASK IBA5-OpcA | Expression of N- terminus Strep-tagged *Synechocystis* OpcA in *E. coli* |
| pASK IBA5-OpcAC410S | Expression of N- terminus Strep-tagged *Synechocystis* OpcAC410S in *E. coli* |
| pASK IBA5-OpcAC192S | Expression of N- terminus Strep-tagged *Synechocystis* OpcAC192S in *E. coli* |
| pASK IBA5-OpcAC192S/C410S | Expression of N- terminus Strep-tagged *Synechocystis* OpcAC192S/C410S in *E. coli* |
| pTK01-Fed9-His | Cloning basis for adding tags |
| pTK30-Fed9-sBit-Spec | Clone cassette containing smBit and Spectamicin resistance |
| pT31Fed9-lBit-Gent | Clone cassette containing lgBit and Gentamicin resistance |
| opcA-Target | Target plasmid containing homolog regions to the c-terminal end of the opcA gene |
| ZWF-Target | Target plasmid containing homolog regions to the c-terminal end of the ZWF (G6PDH) gene |
| opcA-sBit | Tagging opcA with smBit tag in the Genome of Syn6803 |
| opcA-lBit | Tagging opcA with lgBit tag in the Genome of Syn6803 |
| ZWF-sBit | Tagging ZWF with smBit tag in the Genome of Syn6803 |
| ZWF-lBit | Tagging ZWF with lgBit tag in the Genome of Syn6803 |
| pSSRV-sBit | Control plasmid for NanoBit assay |
